# An overexpressed *Q* allele leads to increased spike density and improved processing quality

**DOI:** 10.1101/098558

**Authors:** Bin-Jie Xu, Qing Chen, Ting Zheng, Yun-Feng Jiang, Yuan-Yuan Qiao, Zhen-Ru Guo, Yong-Li Cao, Yan Wang, Ya-Zhou Zhang, Lu-Juan Zong, Jing Zhu, Cai-Hong Liu, Qian-Tao Jiang, Xiu-Jin Lan, Jian Ma, Ji-Rui Wang, You-Liang Zheng, Yu-Ming Wei, Peng-Fei Qi

**Affiliations:** Triticeae Research Institute, Sichuan Agricultural University, Chengdu, Sichuan 611130, China

**Keywords:** Bread-making quality, compact spike, point mutation, protein content, wheat breeding

## Abstract

Spike density and processing quality are important traits during the evolution of wheat, which is controlled by multiple gene loci. The associated gene loci have been heavily studied with slow progress. A common wheat mutant with extremely compact spikes and good processing quality was isolated. The gene (*Q^c1^*) responsible for the mutant phenotype was mapped and cloned, and the cellular mechanism for the mutant phenotype was investigated. *Q^c1^* originated from a point mutation that interferes with the miR172-directed cleavage of the *Q* gene, leading to its overexpression. *Q^c1^* reduces the longitudinal cell size of rachises, resulting in an increased spike density. *Q^c1^* increases the number of vascular bundles, which suggests a higher efficiency in the transportation of assimilates in the spikes of the mutant than in the WT. This could account for the improved processing quality. The effects of *Q^c1^* on spike density and wheat processing quality were confirmed by the identification of nine common wheat mutants having four different *Q^c^* alleles. These results deepen our understanding of the key role of *Q* gene, one of the most important domestication gene for wheat, and provide new insights for the potential application of *Q^c^* allele in wheat breeding.

## Introduction

The spike density of wheat is an important characteristic, which is controlled by multiple gene loci. The expression of the *Compactum* (C) locus on the 2D chromosome of *Triticum aestivum* ssp. *compactum* (Host) Mac Key (club wheat) results in an increased spike density (compact spikes; Dvorak *et al.,* 1998; Johnson *et al.,* 2008). The *Soft outer glume (Sog)* gene located on chromosome 2A^m^ of *Triticum monococcum* (Sood *et al.,* 2009) and the *zeocriton (Zeo)* gene on the 2H chromosome of barley (Druka *et al.,* 2011) can lead to compact spikes as well, and they may be orthologous to *C*. The *Q* gene located on the long arm of chromosome 5A (5AL) affected spike density (Simons *et al.,* 2006). Four gene loci (*C^739^, C^17648^, Cp^m^* and *Cp*) controlling compact spikes have been mapped on 5AL (Mitrofanova, 1997; Laikova *et al.,* 2009; Kosuge *et al.,* 2008, 2012). However, due to the complexity of wheat genome, none of these genes were cloned.

Here, a common wheat mutant (*S-Cp1-1*) with extremely compact spikes was isolated, and the gene responsible for the mutant phenotype was mapped and cloned. A missense mutation in the 10th exon of *Q* reduced its repression by miR172, and this new allele of *Q* gene, *Q^c1^,* dramatically improved the wheat processing quality. The relationship between *Q^c^* alleles (*Q^c1^-Q^c4^*) and the observed phenotype was confirmed by the identification of nine additional mutants. Our results provide new insights for the potential application of *Q^c^* alleles in wheat breeding.

## Materials and Methods

### Wheat materials and growth conditions

The *S-Cp1-1* mutant with compact spikes was isolated from 0.6% EMS-treated common wheat (*Triticum aestivum* L.) cultivar ‘Shumai482’. The WT (*QQ*), heterozygous (*QQ^c1^*) and mutant (*Q^c1^Q*^c1^) lines used in this study were derived from an M6 heterozygous plant. The nine mutants (Supporting information Table S1) were isolated from 0.8% EMS-treated common wheat cultivars ‘Shumai482’, ‘Liangmai4’, ‘Mianmai37’ and ‘Roblin’. To map the *cp1* locus and elevate the effect of *cp1* on the grain protein content (dry weight), we generated two populations from the crosses of *S-Cp1-1* × br220 (a hexaploid wheat line) and *S-Cp1-1* × wanke421 (a common wheat cultivar). The mapping population was grown on the experimental farm of Sichuan Agricultural University in Wenjiang, with a row space of 20 cm x 10 cm. To analyze the processing qualities of WT (*QQ*), heterozygous (*QQ^c^*) and mutant (*Q^c^Q^c^*) lines, the field experiments were performed in a randomized block design with three replicates for each line. Each replicate was planted with an area of 2 m × 4 m, with a row space of 20 cm × 5 cm. A nitrogen: phosphorous: potassium (15: 15: 15; 450 kg per hectare) compound fertilizer was used before sowing.

### Morphological analysis

All of the phenotypic data were analyzed by *X^2^* tests (SPSS 22). The developing spikes of *S-Cp1-1* and WT were scanned using an optical microscope (Olympus, Japan) and EPSON perfection V700 (EPSON, Japan). The spikes at the heading stages were fixed in FAA (70% alcohol: 37% formaldehyde: acetic acid = 18: 1: 1, v: v: v) and embedded in paraffin. Then, the paraffin wax was cut into 6-μm sections using a Leica slicer (Leica, Inc., Germany). Safranin O/ fast green (Solarbio) was used for staining. The splices were photographed using a BX60 light microscope (Olympus, Japan).

### Map-based cloning

Genomic DNA extracted from the young leaves of 260 F2 plants, derived from *S-Cp1-1* × br220, was divided into 24 mixed DNA pools based on their spike phenotypes. The mixed DNA samples were analyzed by Illumina 90K SNP microarray at Compass Biotechnology (Beijing, China), primarily to locate the *cp1* locus. Then, the *cp1* locus was mapped by SSR and STS markers in an F2 population of 819 plants. More molecular markers were developed (Supporting information Table S2) to narrow the candidate region using 10,100 F3 plants derived from heterozygous F2 plants. The BACs and scaffolds were queried using a BLAST algorithm in NCBI (http://www.ncbi.nlm.nih.gov/) and aligned based on their relative positions and overlap. All of the BACs and scaffolds used are listed in Supporting information Table S3.

The cDNA and genomic DNA sequences of candidate genes were amplified from both mutant and WT plants, and confirmed by sequencing (Invitrogen, Shanghai, China). Sequences were analyzed by DNAMAN V6 (Lynnon Biosoft Inc., USA). The promoter of the *Q* gene was amplified with primer pair AP5P.16F + AP5P.12R (Simons *et al.,* 2006) and sequenced.

### Sequence-capture

The qualified genomic DNA sample, extracted by a DNA extraction kit (TIANGEN, Beijing, China), was randomly fragmented. The base-pair peak of the DNA fragments was 200 to 300 bp. Adapters were ligated to both ends of the resulting fragments. DNA was then amplified by ligation-mediated PCR (LM-PCR), purified, and hybridized to the Roche NimbleGen SeqCap EZ Exome probe (BGI, China). Non-hybridized fragments were washed out. Both non-captured and captured LM-PCR products were subjected to quantitative PCR to estimate the magnitude of the enrichment. Each captured library was loaded on a Hiseq4000 platform. We performed high-throughput sequencing for each captured library independently to ensure that each sample met the desired average fold-coverage. Raw image files were processed by Illumina using base calling with default parameters and the sequences of each individual were generated as 100-bp reads. The bioinformatics of sequences were analyzed using BWA (http://bio-bwa.sourceforge.net/) and SAMtools (http://samtools.sourceforge.net/).SOAPsnp (http://soap.genomics.org.cn/soapsnp.html) and SAMtools pileup were used to detect SNPs and InDels, respectively.

### Quantitative RT-PCR analysis

The young spikes of the *S-Cp1-1* mutant and WT lines were collected at three different stages, the pistil and stamen initiation stage, the anther formation stage and the elongation stage. Root, stem and leaf samples of *S-Cp1-1* and WT at the pistil and stamen initiation stage were collected as well. Samples were ground in liquid nitrogen, and RNA was extracted using the Plant RNA extraction kit V1.5 (Biofit, China). Quantitative RT-PCRs were performed using a SYBR premix Ex Taq^tm^ RT-PCR kit (Takara, Dalian, China). All of the experiments were performed following the manufacturer’s instructions. The primers for qRT-PCR are listed in Supporting information Table S2.

### 5’ modified RACE

The 5’ modified RACE was performed as reported previously (Llave *et al.,* 2002). Total RNA was isolated from spikes of the mutant and WT at the pistil initiation stage using a Plant RNA extraction kit V1.5 (Biofit). The primers for the first and second PCR products were Q-cDNA-R and 3’RACE-R (Supporting information Table S2), respectively. The second PCR products were purified and cloned into the pMD19-T vector (Takara, Dalian, China) for sequencing.

### SDS-PAGE analysis

Seed storage proteins were extracted from 20 mg seed powder and separated by sodium dodecyl sulfate-polyacrylamide gel electrophoresis (SDS-PAGE) as described by Qi *et al.* (2011).

### Quantification of proteins by iTRAQ and mass spectrophotometry

The spikes of the mutant and WT at the pistil and stamen initiation stage were collected and ground into powder in liquid nitrogen. The tissue powders were mixed with 200 μL extraction buffer (30 mM Tris-HCl, 2 M thiourea, 7 M urea, 4% 3-[(3-Cholamidopropyl) dimethylammonio] propanesulfonate and 1 mM PMSP inhibitor; pH = 8.5) for 30 min to extract proteins. The total proteins were analyzed by iTRAQ. The molecular weight of the Q^c1^ protein was predicted, and the targeted protein bands in SDS-PAGE were excised from the gel and analyzed by mass spectrophotometry. The BIOWORKS software was used to query, using the BLAST algorithm, the relative proteins in NCBI (Ji *et al.,* 2010). The iTRAQ testing and mass spectrophotometry were conducted at the Beijing Proteome Research Center (Beijing, China).

### Processing quality analysis

Grain samples were cleaned and adjusted to 14% moisture, before being milled with a Brabender Quandrumat Junior mill (Brabender GmbH & Co. KG, Germany). The grain protein content (dry weight), zeleny sedimentation value and wet gluten content were measured following GB/T 17320-2013, using an automatic azotometer (Kjelec 8400; FOSS, Denmark), a zeleny analysis system (CAU-B, China) and a glutomatic 2200 system (Perten, Sweden), respectively.

Dough rheological properties were determined with a 10-g Mixograph (TMCO, Lincoln, USA). Samples were mixed to optimum water absorption following the 54-40A method (AACC, 2001). The development time (min to the curve’s peak) was measured. Finally, results were collected and analyzed using MixSmart software.

The baking test was performed according to AACC method 10.09-01 (AACC, 2010) with some modifications. The baking procedure was the standard rapid-mix-test with 135 g flour at 14% moisture content, and two replicates for each flour sample were performed.

## Results

### *S-Cp1-1* mutant displays a pleiotropic phenotype

A mutant line with spikes as compact as T *compactum* was isolated from ethyl mesylate (EMS)-treated *T. aestivum* cv. ‘Shumai482’, which also shows dwarf characteristics (height) when compared with the wild type (WT). This mutant was named *S-Cp1-1* (the first *Cp* mutant of ‘Shumai482’). The *S-Cp1-1* line had a similar architecture to WT before the tilling stage (Figure 1A). However, by the jointing stage, its plant heights and spike densities were significantly different from those of WT (Figure 1B-1E). A microscopic comparison of the transverse rachises sections revealed that the cells of *S-Cp1-1* were significantly reduced in size and increased in number, and the number of vascular bundles in *S-Cp1-1* was increased (Figure 2A and 2B). Additionally, the vascular morphology was changed in *S-Cp1-1* (Figure 2A). There were lower numbers of xylem cells in the vascular bundles and greater numbers of cells around the vessels (Figure 2A and 2E) when compared with in the WT (Figure 2B and 2F). Longitudinal sections of rachises indicated that the cells in *S-Cp1-1* decreased in size (Figure 2C) compared with in WT (Figure 2D). Backcrossing *S-Cp1-1* with ‘Shumai482’ indicated that the compact spike and dwarf height phenotypes did not segregate in the BC_1_F_2_ population, hiding the fact that these phenotypes in *S-Cp1-1* were controlled by a single locus. We named the locus *compact 1* (*cp1*).

**Figure 1.**
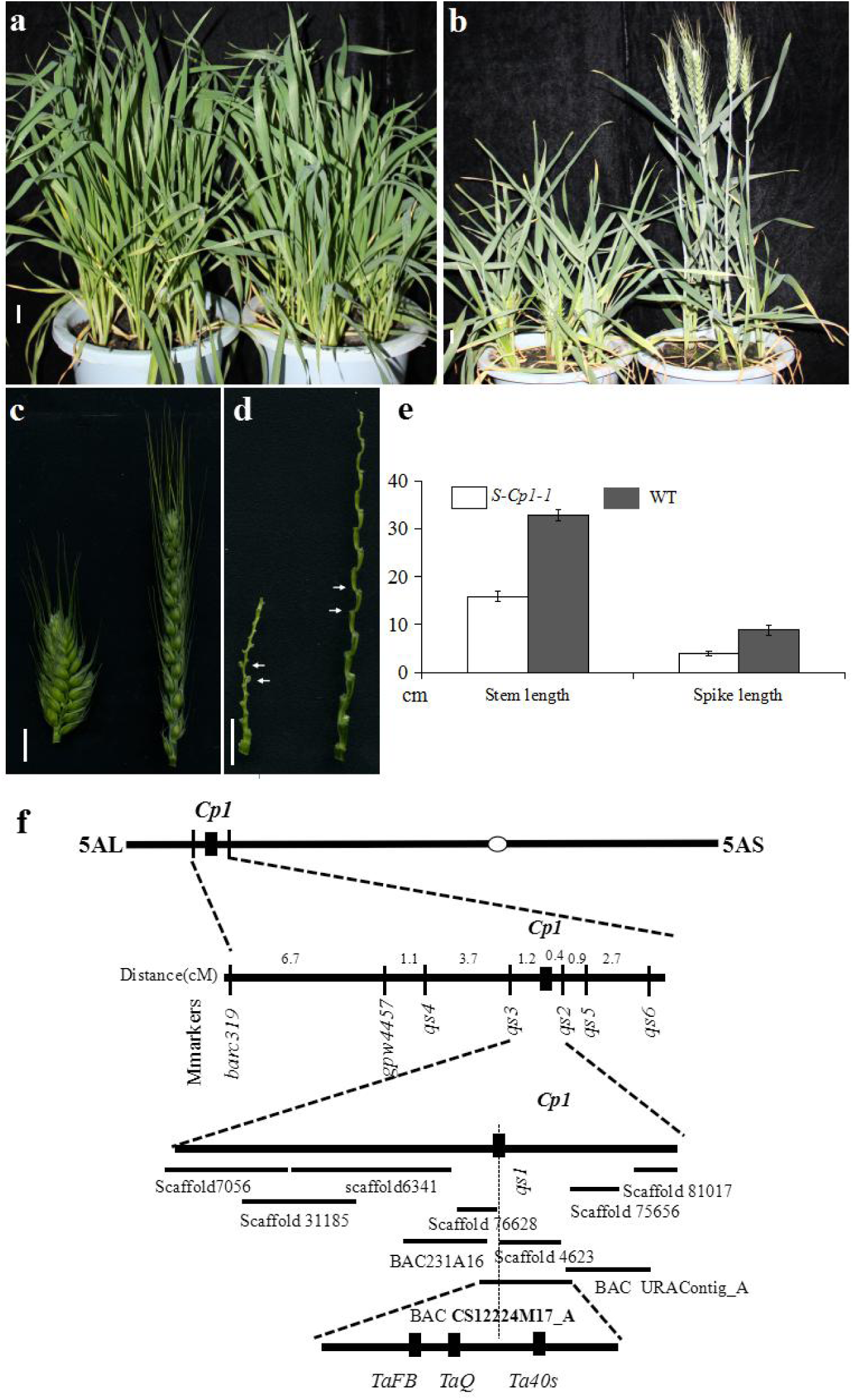
Phenotype of the *S-Cp1-1* mutant and mapping of the *cp1* locus. (A) *S-Cp1-1* (left) and WT (right) plants at the tillering stage. (B) *S-Cp1-1* mutant (left) and WT (right) plants at the heading stage. (C) Spikes of *S-Cp1-1* (left) and WT (right) at the heading stage. (D) Rachises of *S-Cp1-1* (left) and WT (right). The rachis between the white arrows indicates the tissues used in Fig. 2. (E) Stem lengths and spike lengths of *S-Cp1-1* and WT at the maturity stage. Data are means ± s.d. (standard deviation; n = 35). (F) Mapping of the *cp1* locus. Scale bar, 1 cm.

**Figure 2.**
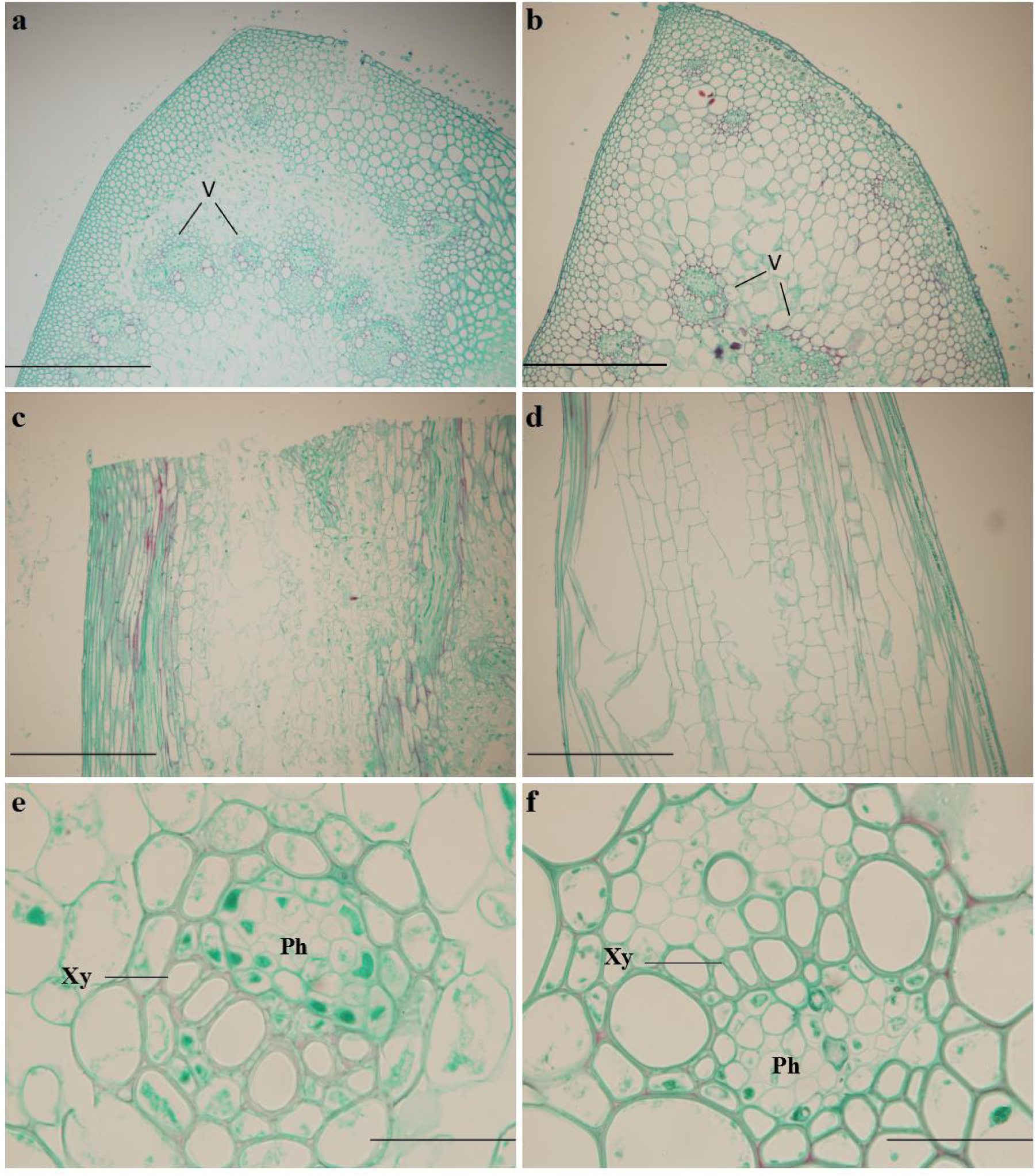
The contrasting cell morphology of the rachises of *S-Cp1-1* and WT. (A)-(B) The transverse sections of *S-Cp1-1* (A) and WT (B). (C)-(D) The longitudinal sections of *S-Cp-1* (C) and WT (D). (E)-(F) The cells in the vascular bundles of *S-Cp1-1* (E) and WT (F). V, vascular bundles; Ph, phloem; Xy, xylem. Scale bar, 10 μm in a-d and 0.1 μm in E and F.

### *Q^c1^* is the candidate gene in the *cp1* locus

A genetic analysis showed that *cp1* was semi-dominant (Supporting information, Figure S1). Compact spikes were selected as the trait for mapping *cp1. cp1* was first positioned on 5AL (Supporting information Figure S2) using a wheat 90 K single-nucleotide polymorphism (SNP) microarray and 24 DNA pools of F2 plants generated from the cross of *S-Cp1-1* and a hexaploid wheat line br220. Furthermore, sequence-tagged site (STS) and simple sequence repeat (SSR) markers (Supporting information Table S2) were developed to place *cp1* in a 1.6-cM region between markers *qs2* and *qs3* with 0.4 cM and 1.2 cM, respectively, based on the physical map draft (Ling *et al.,* 2013; http://plants.ensemble.org;http://www.gramene.org/gremene/searches/ssrtool). More markers were developed to narrow this region, based on the bacterial artificial chromosomes (BACs) and scaffolds in this region (Supporting information Table S3). An STS marker, *qs1*, was identified as co-segregating with *cp1* in 10,100 F_3_ individuals, which were derived from heterozygous F_2_ plants of *S-Cp1-1* × br220. Sequence-capture was performed to sequence the chromosomal region of the WT and *S-Cp1-1* that corresponded to where molecular marker *qs1* was located (GenBank Nos. JF701619 and JF701620, respectively), to identify the possible mutation within this region. A missense mutation (C-T) in the 10th exon of the *Q* gene was found, which was confirmed by gene cloning and sequencing (Genbank Nos. KX580301 and KX580302). This point mutation led to the substitution of serine by leucine (Supporting information Fig. S3). Then, we named the *Q* gene containing this point mutation as *Q^c1^* (the first *Q* with compact spikes).

### Expressional analyses of *Q^c1^*

To determine why *Q^c1^* had an increased spike density, we compared the expression levels of *Q^c1^* and *Q* by qRT-PCR. During spike development, the transcription level of *Q^c1^* was higher than that of *Q* before the anther elongation stage (Figure 3D-3F), and the largest difference was observed at the early spikelet differentiation stages (pistil and stamen initiation stage; Figure 3B-3F). These data indicated that the increased spike density (reduced rachis internodes) in *S-Cp1-1* was a consequence of the higher expression of *Q^c1^* than *Q* (Figure 3A-3D). In addition to differences in spikes, *Q^c1^* and *Q* expressed differentially at the RNA level in roots, stems and leaves, as well as at the pistil and stamen initiation stage. Relative to *Q*, the expression level of *Q^c1^* was much higher in spikes and stems, which was consistent with the phenotypic changes in spike density and plant height (Figure 1B and 1E; Figure 3E).

**Figure 3.**
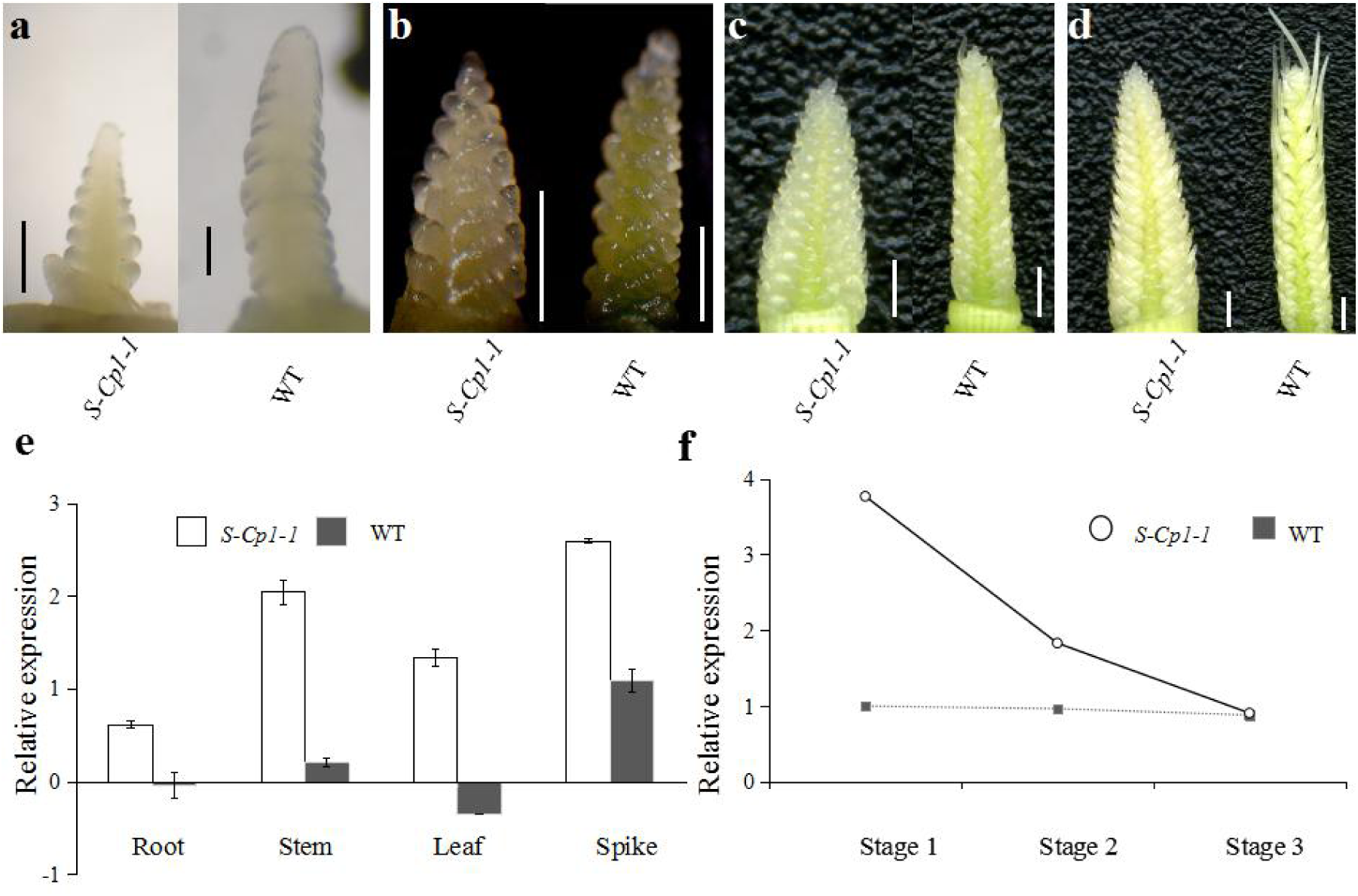
Expressional pattern of *Q*^c1^. (A)-(D) The developing spikes of *S-Cp1-1* (left) and WT (right) at the glume differentiation stage, the pistil and stamen initiation stage, the anther formation stage and the anther elongation stage, respectively. (E) Relative expression levels of *Q^c1^* and *Q* in roots, stems, leaves and spikes at the pistil and stamen initiation stage. (F) Relative expression of *Q^c1^* and *Q* at the pistil and stamen initiation stage (stage 1), the anther formation stage (stage 2) and the anther elongation stage (stage 3). Error bars represent means ± s. ^d^. ^(n^ = ^3)^.

Previous studies demonstrated that *Q* was an AP2 transcription factor member, containing two AP2 DNA-binding domains and a miR172-binding site in the 10th exon (Simons *et al.,* 2006). The *Q^c1^* point mutation was in the miR172-binding site (Figure 4A). To confirm whether the higher expression level of *Q^c1^* was due to the altered cleavage by miR172, a 5’ rapid amplification of cDNA end (RACE) analysis was performed. The sequencing of 20 randomly chosen *Q^c1^* and *Q* clones showed that the cleavage site in the miR172-binding region was changed, resulting in a reduced cleavage efficiency of the miR172 target site. Thus, the point mutation in *Q^c1^* disturbed the *in vivo* cleavage by miR172 (Figure 4B).

**Figure 4.**
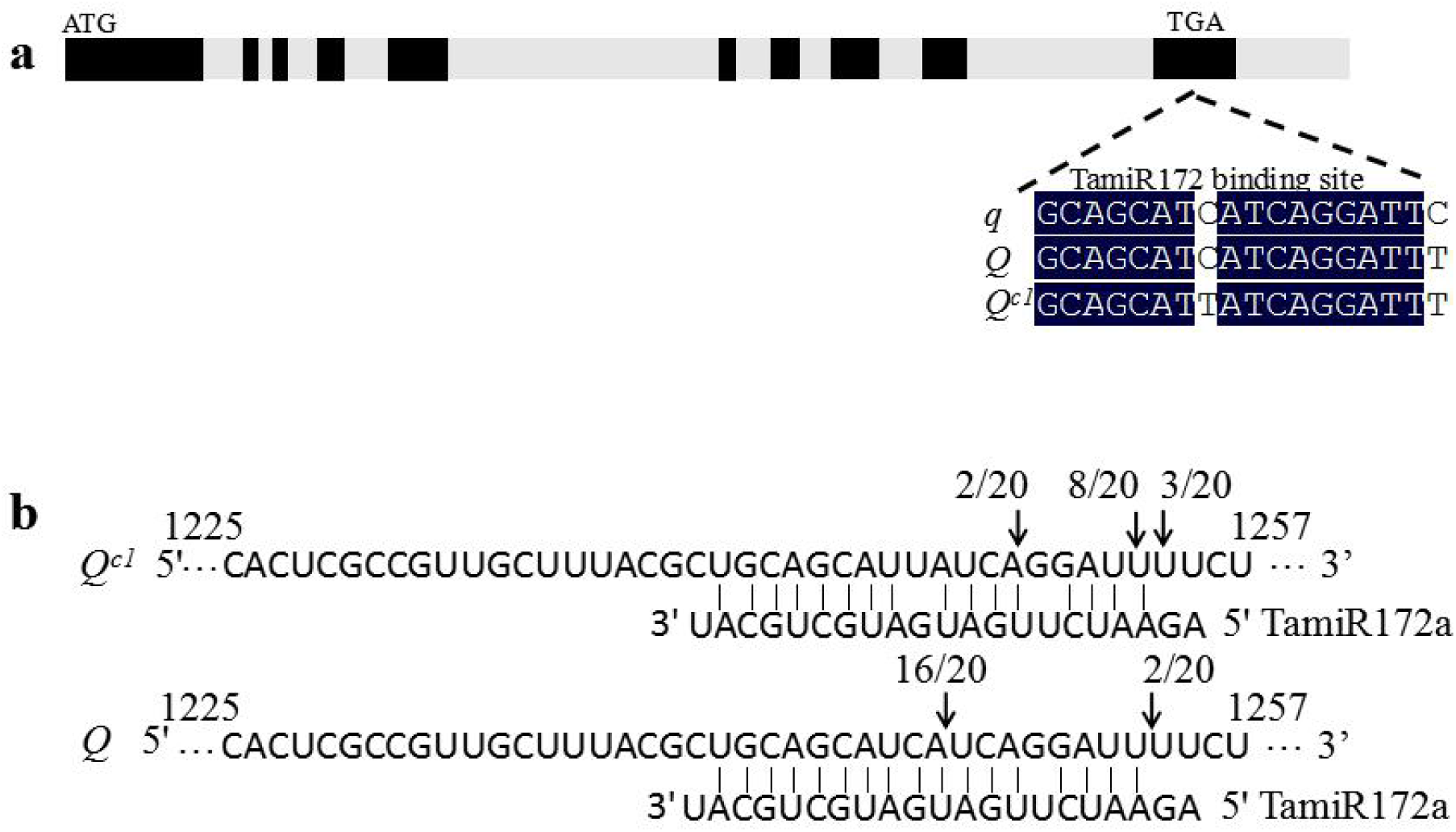
Genomic structure of the *Q^c1^* gene and confirmation of miR172-directed regulation of *Q^c1^.* (A) Genomic structure of the *Q^c1^* gene. The initiation and termination codons, exons (black rectangles) and introns (grey rectangles) are illustrated. The positions of mutations in the miR172-binding region of q, *Q* and *Q^c1^* are indicated. (B) miR172 cleavage sites in the mRNAs of *Q^c1^* and *Q* as determined by RNA ligase-mediated 5’RACE.

### Point mutations in the miR172-binding site of *Q* contributed to the increased spike density

To further confirm the effects of point mutations in the miR172-binding site of the *Q* gene on spike density, common wheat cultivars ‘Shumai482’, ‘Mianmai37’, ‘Liangmai4’ and ‘Roblin’ were treated by EMS again. We obtained, by sequencing, nine independent mutants with four different point mutations in the miR172-binding site of *Q* (Figure 5B; Supporting information Table S1). The spike densities of these nine mutants were similar to that of *S-Cp1-1* (Figure 5A).

**Figure 5.**
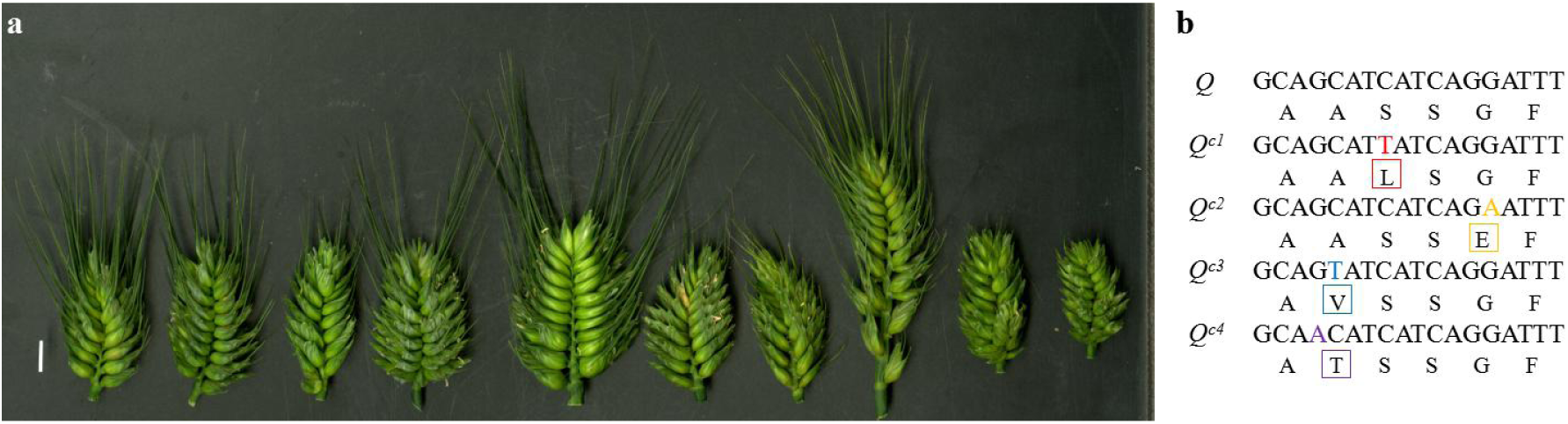
Molecular characterization of *Q^c^* alleles. (A) Features of the spikes of *S-Cp1-1, S-Cp1-2, R-Cp1-3, L-Cp2-1, M-Cp2-2, R-Cp2-3, R-Cp2-4, S-Cp3-1, R-Cp3-2* and *R-Cp4-1* (left to right; Supporting information Table S1). (B) Polymorphisms of the *Q^c^* alleles and their predicted amino acid substitutions.

### *Q^c^* alleles improve the wheat processing quality

The increased number of vascular bundles in *S-Cp1-1* (Figure 2) suggested a higher transportation efficiency of assimilates in the spikes of mutants than in those of WT, which motivated us to compare the processing qualities of *S-Cp1-1* and WT. Dramatic differences were found when comparing the processing parameters of the *S-Cp1-1* mutant, heterozygote and WT lines (Table 1), and the loaf volume of the mutant line was much greater than that of the WT (Supporting information Figure S4). Notably, no variations in seed storage proteins were observed (Supporting information Figure S5), especially the high molecular weight glutenin subunits, which are among the most important determinants in bread-making (Shewry *et al.,* 2003). To demonstrate that the improvement in the wheat processing quality was due to the presence of *Q^c1^*, the grain protein contents (dry weight) of two F2 populations were measured. As expected, the grain protein contents of plants with two copies of the *Q^c1^* gene were much higher than those with one or no *Q^c1^* copies (Figure 6). The positive effect of *Q^c^* alleles on the processing quality was supported by comparing the grain protein contents of mutants having different *Q^c^* alleles to the WT (Supporting information Table S1).

**Table 1.** Effect of *Q^c1^* on wheat processing quality parameters.

| Genotype | Grain protein content | Wet gluten | Sedimentation | Development |
| --- | --- | --- | --- | --- |
|  | (%; dry weight) | content (%) | value (mL) | time (min) |
| $Q^{c1}/Q^{c1}$ | 19.73a | 50.60a | 63.05a | 7.23a |
| $Q^{c1}/Q$ | 16.20b | 39.63b | 50.52b | 5.42b |
| $Q/Q$ | 14.00c | 34.83c | 36.83c | 3.55c |

Different letters represent significance at $P < 0.05$ . Significance was calculated using Student's $t$ -test (DPS software version 12.01; Tang & Zhang 2013). The WT ( $QQ$ ), heterozygous ( $QQ^c$ ) and mutant ( $Q^cQ^c$ ) lines used were derived from an $M_6$ heterozygous plant.

**Figure 6.**
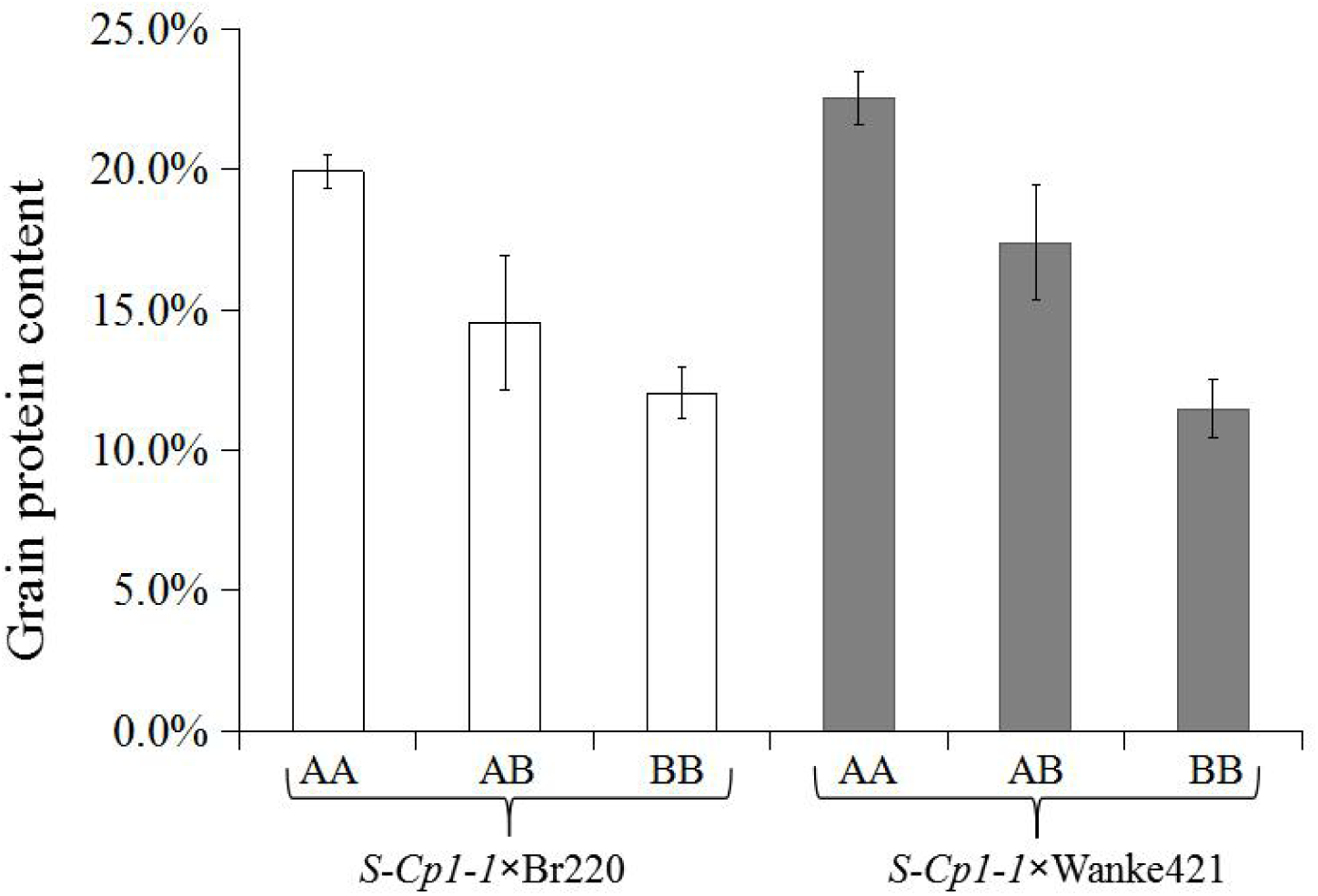
Grain protein contents of seeds harvested from individual plants in two F2 populations. Data are means ± s.d. (n = 20). The different small letters above each box indicate significance at P < 0.05.

## Discussion

Numerous studies have explored the genes determining and affecting spike density in wheat. It was believed that the dominant allele of the *C* gene determined the compact spike phenotype of club wheat, which was located on chromosome 2D (Johnson *et al.,* 2008). However, research on four compact determining genes (*C^17648^, C^739^, Cp* and *Cp^m^*) from wheat mutants indicated that a locus in 5AL also affected the compact spike phenotype (Mitrofanova, 1997; Laikova *et al.,* 2009; Kosuge *et al.,* 2008, 2012). Further, it was hypothesized that the genomic region affecting spike density flanked the *Q* locus (Kosuge *et al.,* 2012). Simons *et al.* (2006) reported that a transgene of *Q* in bread wheat could occasionally lead to compact spikes. Our research illustrated that the *Q* gene with point mutations in the miR172-binding region determined the spike density of wheat.

For miRNA-directed cleavage, base-pairing between miRNAs and their target mRNAs is critical (Huntzinger and Izaurralde, 2011). The point mutations in the miRNA-binding region of the *Q* gene interfered with the regulation of miR172 and thus led to an increased spike density and decreased plant height. *Q* was one of the most important genes during the domestication of bread wheat. A single point mutation in the 10th exon of *q* results in the emergence of Q, which caused desired changes in wheat, including the loss of glume and rachis fragility (Simons *et al.,* 2006). The point mutation in *q* did not result in an amino acid difference. This indicates that the compact spike phenotype is mainly determined by the disturbance in the cleavage of miR172 rather than a protein sequence change.

miR172 mainly regulates its target mRNAs by translational inhibition and/or transcript cleavage in Arabidopsis (Chen, 2004). However, the expressional difference at the RNA level was easily observed (Figure 3) when comparing *S-Cp1-1* and WT. To determine whether there was any difference at the protein level, isobaric tags for relative and absolute quantification (iTRAQ) and mass spectrophotometry were carried out. Unfortunately, the Q protein was not detected by these two methods, possibly because of the special structure of Q and the limitations of the techniques.

Plant height is an important agronomic trait for plant architecture and grain yield in wheat. *Reduced-height 1 (Rht1)*, which gave rise to the “green revolution”, and other *Rht* genes have been used successfully in wheat breeding (Peng *et al.,* 1999; Pearce *et al.,* 2011; Chen *et al.,* 2014). Most *Rht* genes control the dwarf phenotype by regulating endogenous phytohormones (Hong *et al.,* 2003; Tanabe *et al.,* 2005). Additionally, the phenylpropanoid pathway is involved in the dwarf phenotype (Schoor *et al.,* 2011). The silencing of the *cinnamyl alcohol dehydrogenase* gene could lead to a dwarf phenotype with a reduced lignin content (Trabucco *et al.,* 2013). Because of the reduced lignin content in the culm of *S-Cp1-1* compared with WT (data not shown), we speculated that the phenylpropanoid pathway in *S-Cp1-1* was affected by the higher expression of *Q^c1^* in the stem, which resulted in the decreased plant height in the mutant.

The improvement in the wheat processing quality has been extensively studied over the years. Grain protein content is a crucial index for measuring wheat quality (Weegels *et al.,* 1996). *S-Cp1-1* had a higher grain protein content than WT, suggesting a key role of *Q^c1^* in regulating the accumulation of seed storage proteins in wheat. A transcriptome analysis (unpublished data) showed that the expression of the *storage protein activator* gene in *S-Cp1-1* increased by 7-fold, compared with WT. Therefore, we suspected that the overexpression of *Q* promoted the expression of the gene and subsequently enhanced the biosynthesis of seed storage proteins (Ravel *et al.,* 2009). The significant change in the processing quality in common wheat mutants with different *Q^c^* alleles indicated that *Q^c^* can be used in quality breeding.

### Accession Numbers

Nucleotide sequence data from this article can be found in the GenBank/EMBL databases under the following accession numbers:KX580301-KX580304 and KX620761-KX620768.

## Author contributions

PF.Q. and Y.L.Z. designed the experiments. P.F.Q., B.J.X., Q.C., Z.T., Y.M.W., J.R.W., Q.T.J., X.J.L., J.M. and Y.L.Z. analyzed the data. P.F.Q. and Q.C. prepared the plant materials. P.F.Q., B.J.X. and Q.C. wrote the manuscript. P.F.Q., B.J.X., Y.Z.Z., Y.L.Z. and Y.M.W. prepared the figures. P.F.Q. , B . J . X. , Q . C . , T.Z . , Y.F. J. , Y Y Q . , Z R. G. , Y.L . C . , Y.W., Y.Z.Z., L.J.Z. J.Z. and C.H.L. performed the experiments. J.R.W., Q.T.J., X.J.L. and J.M. provided key advice.

## Acknowledgements

This research was supported by the National Natural Science Foundation of China (31230053, 31570335 and 31671677), and the National Basic Research Program of China (2014CB147200). We thank Dr. Hong-Yang Yu, Dr. Jing Fan and Prof. Wen-Ming Wang at Sichuan Agricultural University for their technical assistance.

## Supplemental files

**Figure S1** Phenotypes of *Q^c1^/Q^c1^, Q^c1^/Q, Q/Q* in the common wheat cultivar ‘Shumai482’ background. The spike density and plant height of the heterozygote were intermediate values.

**Figure S2** Mapping of the *cp1* locus using SNP markers. Dots indicate the markers. The ordinate represents the square of the correlation coefficient.

**Figure S3** Alignment of the amino acid sequences of the *Q* and *Q^c^* alleles. The conserved domains are indicated (Gil-Humanes *et al.,* 2009).

**Figure S4** Effects of *Q^c1^* on loaf volume.

**Figure S5** Separation of seed storage proteins by SDS-PAGE. HMW-GS, high molecular weight glutenin subunit; LMW-GS, low molecular weight glutenin subunit; Ax1, Dx5, Bx7, By9 and Dy10 indicate the composition of HMW-GS in the wheat cultivar “Shumai482”.

**Table S1** The effects of four *Q^c^* alleles on the grain protein content (dry weight).

**Table S2** Primers used in this study.

**Table S3** Scaffolds and BACs used in this study.

## References

AACC., 2001 Sedimentation test for wheat. Approved Methods of the American Association of Cereal Chemists (Method 56-61), 10th ed. AACC, St. Paul, MN, USA

AACC., 2010 Basic Straight-Dough Bread-baking Method. Approved Methods of the American Association of Cereal Chemists (Method 10.09-01), St. Paul, MN, USA

Chen, L., L. Hao, A. G. Condon, and Y. G. Yu, 2014 Exogenous GA3 application can compensate the morphogenetic effects of the GA-responsive dwarfing gene *Rht12* in bread wheat. Plos One 9: e86431.

Chen, X., 2004 A microRNA as a translational repressor of *APETALA2* in *Arabidopsis* flower development. Science 303: 2022–2025.

Dvorak, J., M. C. Luo, Z.L. Yang, and H.B. Zhang, 1998 The structure of the *Aegilops tauschii* genepool and the evolution of hexaploid wheat. Theor. Appl. Genet. 97: 657–670.

Druka, A., J. Franckowiak, U. Lundqvist, N. Bonar, J. Alexander et al., 2011 Genetic dissection of barley morphology and development. Plant Physiol. 155: 617–627.

Faris, J. D., J. P. Fellers, S. A. Brooks, and B. S. Gill, 2003 A bacterial artificial chromosome contig spanning the major domestication locus *Q* in wheat and identification of a candidate gene. Genetics 164: 311–321.

Faris, J. D., Z. Zhang, J. P. Fellers, and B. S. Gill, 2008 Micro-colinearity between rice, *Brachypodium,* and *Triticum monococcum* at the wheat domestication locus Q. Funct. Integr. Genomics 8: 149–164.

Gil-Humanes, J., F. Pistón, A. Martín, and F. Barro, 2009 Comparative genomic analysis and expression of the *APETALA2-like* genes from barley, wheat, and barley-wheat amphiploids. BMC Plant Biol. 9: 66.

Hong, Z., M. Ueguchi-Tanaka, K. Umemura, S. Uozu, S. Fujioka, 2003 A rice brassinosteroid-defective mutant, *ebisu dwarf* (d2), is caused by a loss of function of a new member of cytochrome P450. Plant Cell 17: 776–790.

Huntzinger, E., and E. Izaurralde, 2011 Gene silencing by microRNAs: contributions of translational repression and mRNA decay. Nat. Rev. Genet. 12: 99–110.

Ji, L., T. Barrett, O. Ayanbule, D. B. Troup, D. Rudnev, 2010 NCBI Peptidome: a new repository for mass spectrometry proteomics data. Nucleic Acids. Res. 38: D731–D735.

Johnson, E. R., V. J. Nalam, R. S. Zemetra, and O. Riera-Lizarazu, 2008 Mapping the compactum locus in wheat (*Triticum aestivum* L.) and its relationship to other spike morphology genes of the *Triticeae*. Euphytica 163: 193–201.

Kosuge, K., N. Watanabe, T. Kuboyama, V. M. Melnik, V. I. Yanchenko, 2008 Cytological and microsatellite mapping of mutant genes for spherical grain and compact spikes in durum wheat. Euphytica 159: 289–296.

Kosuge, K., N. Watanabe, V. M. Melnik, L. I. Laikova, and N. P. Goncharov, 2012 New sources of compact spike morphology determined by the genes on the chromosome 5A in hexaploid wheat. Genet. Resour. Crop Evol. 59: 1115–1124.

Laikova, L. I., N. P. Goncharov, O. P. Popova, V. M. Melnik, O. P. Mitrofanova, and N. Watanabe, 2009 Genetic studies of bread wheat mutants. Bull. Appl. Bot. Genet. Breed 166: 396–399.

Ling, H. Q., S. Zhao, D. Liu, J. Wang, H. Sun, et al., 2013 Draft genome of the wheat A-genome progenitor *Triticum urartu*. Nature 496: 87–90.

Llave, C., Z. Xie,K. D. Kasschau, and J. C. Carrington, 2002 Cleavage of Scarecrow-like mRNA targets directed by a class of *Arabidopsis* miRNA. Science 297: 2053–2056.

Mitrofanova, O. P., 1997 The inheritance and effect of *Cp* (Compact plant) mutation induced in common wheat. Genetika 33: 482–488.

Pearce, S., R. Saville, S. P. Vaughan, P. M. Chandler, E. P. Wilhelm, et al., 2011 Molecular characterization of *Rht-1* dwarfing genes in hexaploid wheat. Plant Physiol. 157: 1820–1831.

Peng, J., D. E. Richards, N. M., Hartley, G. P. Murphy, K.M. Devos, et al., 1999 “Green revolution” genes encodes mutant gibberellin response modulators. Nature 400: 256–261.

Qi, P. F., Y. M. Wei, Q. Chen, T. Quellet, J. Ai, G. Y. Chen, W. Li, Y. L. Zheng, 2011 Identification of novel α-gliadin genes. Genome 54: 244–252.

Ravel, C., P. Martre, I. Romeuf, M. Dardevet, R. El-Malki, et al, 2009 Nucleotide polymorphism in the wheat transcriptional activator *Spa* influences its pattern of expression and has pleiotropic effects on grain protein composition, dough viscoelasticity, and grain hardness. Plant Physiol. 151: 2133–2144.

Schoor, S., S. Farrow, H. Blaschke, S. Lee, G. Perry, et al., 2011 Adenosine kinase contributes to cytokinin interconversion in *Arabidopsis*. Plant Physiol. 157: 659–672.

Schwab, R., J. F. Palatnik, M. Riester, C. Schommer, M. Schmid, et al., 2005 Specific effects of microRNAs on the plant transcriptome. Dev. Cell 8: 517–527.

Shewry, P. R., N. G. Halford, A. S. Tatham, Y. Popineau, D. Lafiandra, et al., 2003 The high molecular weight subunits of wheat glutenin and their role in determining wheat processing properties. Adv. Food Nutr. Res. 45: 219–302.

Simons, K. J., J. P. Fellers, H. N. Trick, Z. Zhang, Y. Tai, et al., 2006 Molecular characterization of the major wheat domestication gene Q. Genetics 172: 547–555.

Sood, S., V. Kuraparthy, G. Bai, B. S. Gill, 2009 The major threshability genes soft glume *(sog)* and tenacious glume (Tg), of diploid and polyploid wheat, trace their origin to independent mutations at non-orthologous loci. Theor. Appl. Genet. 119: 341–351.

Tang, Q. Y., and C. X. Zhang, 2013 Data Processing System (DPS) software with experimental design, statistical analysis and data mining developed for use in entomological research. Insect Sci. 20: 254–260.

Tanabe, S., M. Ashikari, S. Fujioka, S. Takatsuto, S. Yoshida, et al., 2005 A novel cytochrome P450 is implicated in brassinosteroid biosynthesis via the characterization of a rice dwarf mutant, *dwarf11,* with reduced seed length. Plant Cell 17: 776–790.

Trabucco, G. M., D. A. Matos, S. J. Lee, A. J. Saathoff, H. D. Priest, et al., 2013 Functional characterization of cinnamyl alcohol dehydrogenase and caffeic acid O-methyltransferase in Brachypodium distachyon. BMCBiotechnol. 13:61

Wang, S., D. Wong, K. Forrest, A. Allen, S. Chao, et al., 2014 Characterization of polyploid wheat genomic diversity using a high-density 90, 000 single nucleotide polymorphism array. Plant Biotechnol. J. 12: 787–796.

Weegels, P. L., R. J. Hamer, and J. D. Schofield, 1996 Critical review: functional properties of wheat glutenin. J. Cereal Sci. 23: 1–18.

Yan, L., A. Loukoianov, G. Tranquilli, M. Helquera, T. Fahima, et al., 2003 Positional cloning of the wheat vernalization gene of *VRN1*. Proc. Natl. Acad. Sci., USA 100: 6253–6268.

Zhang, Z., H. Beclam, P. Gornicki, M. Charlesb, J. Justb, et al., 2011 Duplication and partitioning in evolution and function of homoeologous *Q* loci governing domestication characters in polyploid wheat. Proc. Natl. Acad. Sci., USA 108: 18737–18742.

